# Optimistic ants: Positive cognitive judgement bias but no emotional contagion in the ant *Lasius niger*

**DOI:** 10.1101/2022.11.03.515024

**Authors:** K. Wenig, H. Kapfinger, A. Koch, T.J. Czaczkes

## Abstract

Understanding the emotional states of animals is key for informing their ethical treatment, but very little attention has been directed towards the emotional lives of invertebrates. As emotions influence information processing, one way to assess emotional states is to look for an individual’s cognitive bias, i.e., their tendency to make optimistic or pessimistic judgements. Here we developed a free-running judgment bias task for the ant *Lasius niger*, and applied the judgement bias to assess ants’ reactions towards positive and negative stimuli. After an initial learning phase in which individuals were trained to associate two odour stimuli with positive or negative reinforcement, their reaction towards ambiguous stimuli, i.e., a mixture between both odours, was assessed. We also explored our study species’ capacity to socially transmit emotional states (‘emotional contagion’) by investigating whether social information could elicit emotional responses. We find *L. niger* to be optimistic, showing a baseline positive judgement bias, with 65-68% of ants preferring an ambiguous 1:1 mix of positive and negative cues over no cues. Providing an unexpected food reward prior to the judgement bias task increases positive judgement bias (c. 75% positive). There was a non-significant tendency towards a negative judgement bias after experiencing a mild electric shock (c. 75% negative). Neither positive nor negative social information (trail and alarm pheromones, respectively) affected the ants’ judgement biases, thus providing no indication for emotional contagion. The development of a powerful, simple, and ecologically relevant cognitive judgement task, deployable in the lab and in the field, opens the door to systematic comparative studies of the evolutionary and ecological causes of judgement bias.

## Introduction

> “*It’s ok to eat fish*
>
> *because they don’t have any feelings”*
>
> from *‘Something in the Way’, Nirvana, 1991*

Having subjective emotions is considered by many as a key criterion for protecting an entities’ well-being. Emotions are short-lived and event-related processes encompassing subjective, behavioral, physiological, and cognitive components (Mendl, Oliver & Paul, 2010). Due to the inability to self-report on the subjective component, the emotional states of non-human animals’ can only be assessed via behavioral changes, e.g., in locomotor activity (Moe et al., 2006), vocalizations (Knutson, 2002), self-directed (Maestripieri et al., 1992), affiliative (Clegg et al., 2017), exploratory, and play behaviours (Murphy et al., 2014; Zupan et al., 2016); or via physiological changes, e.g., in corticosteroid levels (M. A. Novak et al., 2013), body surface temperature (Robinson et al., 2012), or heart rate (Von Borell et al., 2007).

Cognitive components can be evaluated by measuring the effect of emotional experiences on information processing functions, such as memory (Burman & Mendl, 2018), attention (Bethell et al., 2016; Crump et al., 2018), or appraisal. An important test for evaluating emotional experiences is the non-invasive judgement bias task. Here, individuals are trained on two cues, one of which predicts a positive event (e.g., highly preferred food) while the other predicts a less positive (e.g., less preferred food) or negative event (e.g., negative reinforcement). After training, ambiguous cues can be used to test the subjects’ judgment biases: subjects in a putative positive emotional state are expected to express a tendency to categorize ambiguous cues as predictive of a positive outcome (‘optimistic/positive judgment bias’) while individuals in a supposedly negative emotional state are anticipated to categorize ambiguous cues as predictors for negative outcomes (‘pessimistic/negative judgment bias’, (Baciadonna & McElligott, 2015; Mendl et al., 2009; Paul et al., 2005).

The judgement bias task has become increasingly important for the study of animal welfare, and has been applied to various species (see Lagisz et al., 2020 for a review), predominantly in captive settings (Bethell, 2015): Farm livestock (Baciadonna & McElligott, 2015), horses (Henry et al., 2017), non-human primates (Bateson & Nettle, 2015; Bethell et al., 2012), dolphins (Clegg et al., 2017), dogs (Burman et al., 2011; Mendl, Brooks, et al., 2010), rodents (Bethell & Koyama, 2015; Nguyen et al., 2020), songbirds (Bateson & Matheson, 2007; Brilot et al., 2009; Matheson et al., 2008; McCoy et al., 2019), fish (Tan, 2017) and also, although very rarely, insects. Honeybees (*Apis mellifera carnica*) and fruit flies (*Drosophila melanogaster*) were found to more likely classify ambiguous cues as predictive of a negative outcome after they encountered an artificial predator attack, simulated by vigorous shaking (Bateson et al., 2011; Deakin et al., 2018; Schlüns et al., 2017). Bumblebees (*Bombus terrestris*) that received an unexpected reward (droplet of sucrose solution) expressed optimism in a judgment bias task and subsequently also recovered faster from a experimentally induced predator attack (Solvi et al., 2016). Carpenter ants showed individual differences in the judgment bias task, with fast explorers being more likely to express a pessimistic bias and slow explorers more likely to show an optimistic bias (d’Ettorre et al., 2017). Importantly, however, Baracchi et al. (2017) challenged the interpretation of these results (mainly those of Solvi et al., 2016) by suggesting alternative, arguably more parsimonious explanations of the observed behaviors (an increase in appetitive motivation due to an unexpected sucrose reward rather than an optimistic internal, emotion-like state) and overall urges caution when adapting anthropomorphically biased terminology to invertebrates.

From a welfare point of view, it is not only the emotional state of a single individual which is relevant, but also the extent to which individuals are affected by their conspecifics’ emotional expressions (Baciadonna et al., 2018; Düpjan, 2020). For example, pigs that were confronted with positively or negatively manipulated pen mates consequently showed carry- over effects on their own emotional states (Reimert et al., 2017). The process of transmission of emotional states between individuals is called ‘emotional contagion’ and is suggested to facilitate coordination and communication in social groups (Adriaense et al., 2020; Hatfield et al., 1993). While the judgement bias task offers a valuable opportunity to assess emotional contagion, only few studies have used this cognitive approach so far: In ravens, manipulated individuals and unmanipulated observers expressed a negative, but not a positive, judgment bias after respective emotional manipulations (Adriaense et al., 2019). Rats showed both positive and negative emotional contagion, measured via the judgement bias task, after hearing recordings of positive (50 kHz) or negative (22 kHz) ultrasonic vocalizations (Saito et al., 2016).

Many eusocial insects, such as ants and bees, live in social groups that show a high degree of coordination, with individuals communicating via visual (Dyer, 2002; Michelsen, 2003), mechanical (Hunt & Richard, 2013), and especially chemical information (Orlova, 2019; von Thienen et al., 2014). However, their emotional lives are largely disregarded, as they are often considered to be as a ‘lower class’ or ‘simpler’ in comparison to vertebrates (Baracchi et al., 2017; Mikhalevich & Powell, 2020; Perry et al., 2017; Perry & Baciadonna, 2017). While the emotional states of invertebrates in general have just begun to be explored (Perry & Baciadonna, 2017), the transmission of emotional states in socially-living invertebrate species has not, to our knowledge, been investigated so far.

Here, we developed a straightforward and ecologically relevant free-running assay for testing judgement bias in the black garden ant (*Lasius niger*). In a learning phase, we carried out a standard alternative discrimination training with a positive reinforcement associated to one odour (‘positive odour’) and a negative reinforcement associated to another odour (‘negative odour’). The preference of the trained ants to various odours is then tested on a Y-maze. In a first experiment, we used mixtures of positive and negative odours in different ratios to determine a suitable ambiguous odour stimulus for our study species, i.e., the odour mixture that ants chose over an unscented control option in about 50% of trials. Then, in a second experiment, we applied specific treatments in order to manipulate optimism and pessimism: Before entering the judgement bias task, ants either encountered i.) a 3-volt electric shock, ii.) a tiny drop of sucrose solution, iii.) alarm pheromones, indicating danger or iv.) trail pheromones, indicating a food resource. We expected positively manipulated subjects (sucrose and trail pheromone treatment) to exhibit an optimistic judgement bias while we expected negatively manipulated subjects (electric shock and alarm pheromone treatment) to categorize the ambiguous odour as predictive of a negative outcome. While the sucrose and electric shock treatments used direct experience (=private information), the alarm and trail pheromone treatments represented social information, which we used to test whether optimism and pessimism can be socially transmitted via emotional contagion.

## Methods

### Study species and maintenance

16 queenless colony fragments (henceforth ‘colonies’) of the black garden ant, *Lasius niger*, were gathered from 16 different mother colonies on the University of Regensburg campus. Ant colonies were housed in plastic boxes (40 × 30 × 20 cm) containing a circular plaster nest (14 cm diameter, 2 cm high) and with a layer of plaster covering the bottom. Each colony comprised around 500–1500 workers and small amounts of brood. Queenless colonies forage and lay pheromone trails, and are frequently used in foraging experiments (Detrain et al., 2019) as due to the rare interactions of foragers with the queen (Stroeymeyt et al., 2018), a lack of queen (but not brood; see Portha et al., 2004) should have little effect on the foragers’ behaviour. Colonies were fed *ad libitum* on 0.5 M sucrose solution and received *Drosophila melanogaster* fruit flies. Four days prior to the experiment, colonies were deprived of food in order to achieve a uniform and high motivation for foraging (Josens & Roces, 2000; Mailleux et al., 2006). Water was always available *ad libitum*. Colonies were kept at 25 ° C (+/-2) with 12-hour photoperiodic cycle.

### Experimental setups and procedures

#### Experiment 1: Developing a free-running judgement bias task for ants

##### a)Testing different odour ratios

In a learning phase, we carried out a standard alternative discrimination training on a linear runway. Ants were trained to associate one odour with a positive reinforcement (‘positive odour’) and another odour with a negative reinforcement (‘negative odour’). As *L. niger* learned the association within a single trial in comparable experiments (Wenig et al., 2021), here we also confronted them only one time each with the positive and negative odours. To start the experiment, a drawbridge was lowered, giving access to the setup. The first ant to climb onto the bridge was allowed to proceed to a 20 cm linear runway which was scented with lemon or rose odour. The positive association trial was always presented first to ensure the subject’s motivation for participation. After reaching a drop of 1M sucrose solution at the end of the runway, flavoured to match the runway odour, the ant was marked with acrylic paint on its abdomen while drinking. After drinking to satiation, she was allowed to return the nest. After unloading the sugar, the ant was brought back for the negative association trial using a drop of bitter-tasting quinine solution (60mM) – the procedure was the same as with the positive reinforcement, but if the ant did not return to the nest after tasting the quinine solution, she was gently returned to the nest box with a piece of paper. Lemon and rose odours were used, balanced across subjects between having a positive and negative association. We scented runway paper layers by placing unscented layers in an airtight container with a glass petri dish containing 500 μL of food flavouring for at least 10 hours. To intensify the association, we also added rose and lemon food flavouring to the respective sucrose solution and quinine drop (ratio: 1 μL per mL).

In the test phase, we prepared a Y-shaped setup (‘Y-maze’) with a stem and two arms (10cm long, 1cm wide, tapering to 2mm at the bifurcation). The stem and one arm were covered with unscented paper overlays, while the other arm was covered with a scented paper overlay. The scented overlays were scented with rose and lemon odours of different ratios: We combined the previously positively and negatively associated scents in ratios of 3:1, 1:1, 1:3, 1:6 and 1:9, respectively. Scenting was achieved as above but using a mix of both food flavourings. The ant was considered to have made a decision when it crossed a line 2cm from the Y-maze bifurcation.

After the ants’ preference was tested, we ran a memory probe to ensure that learning had, indeed, taken place. Subjects were transferred to a second Y-maze, with one arm carrying the positively associated odour and the other arm covered with the negatively associated odour. Again, we recorded the subject’s decision when running at least 2 cm on one of the Y-maze arms.

In total, 275 ants were tested (3:1 ratio: n=26; 1:1 ratio: n=32; 1:3 ratio: n=24; 1:6 ratio: n=96, 1:9 ratio: n=97).

##### b) Increased and unavoidable negative reinforcement

A striking finding from experiment 1a was that ants show a high baseline positive judgement bias (see results). We feared that this may have been due to either the quinine solution alone being a too-weak negative stimulus, or because ants can in fact easily avoid negative quinine reinforcement by carefully probing solution drops, and rejecting bad-tasting ones in the future (Wenig et al., 2021). To test this, we repeated the 1:1 ratio experiment but increased negative reinforcement by adding an electric shock apparatus on the negatively reinforced runway, 1cm from the quinine drop, to increase the cost and make the negative reinforcement unavoidable. The electric shocker was built by affixing two copper strips next to each other with a narrow gap (c. 0.5mm) to ensure that ants touched both strips simultaneously when walking across the copper strips, thus closing the electric circuit. The copper strips were connected to a nine-volt battery. For positive and negative associations rose and orange odours were used, balanced across subjects. In total, 65 ants were tested.

#### Experiment 2: Judgement biases

Based on the results of the first experiment, we used a 1:6 ratio mixture of positive and negative odours to measure ants’ judgement biases. Note that while a ratio of 1:9 may have been slightly more appropriate (see figure 1), due to experimental error 1:6 was used. However, as ants were also indifferent between this mix ratio and an unscented overlay (see results), this should not influence the results.

**Fig. 1:**
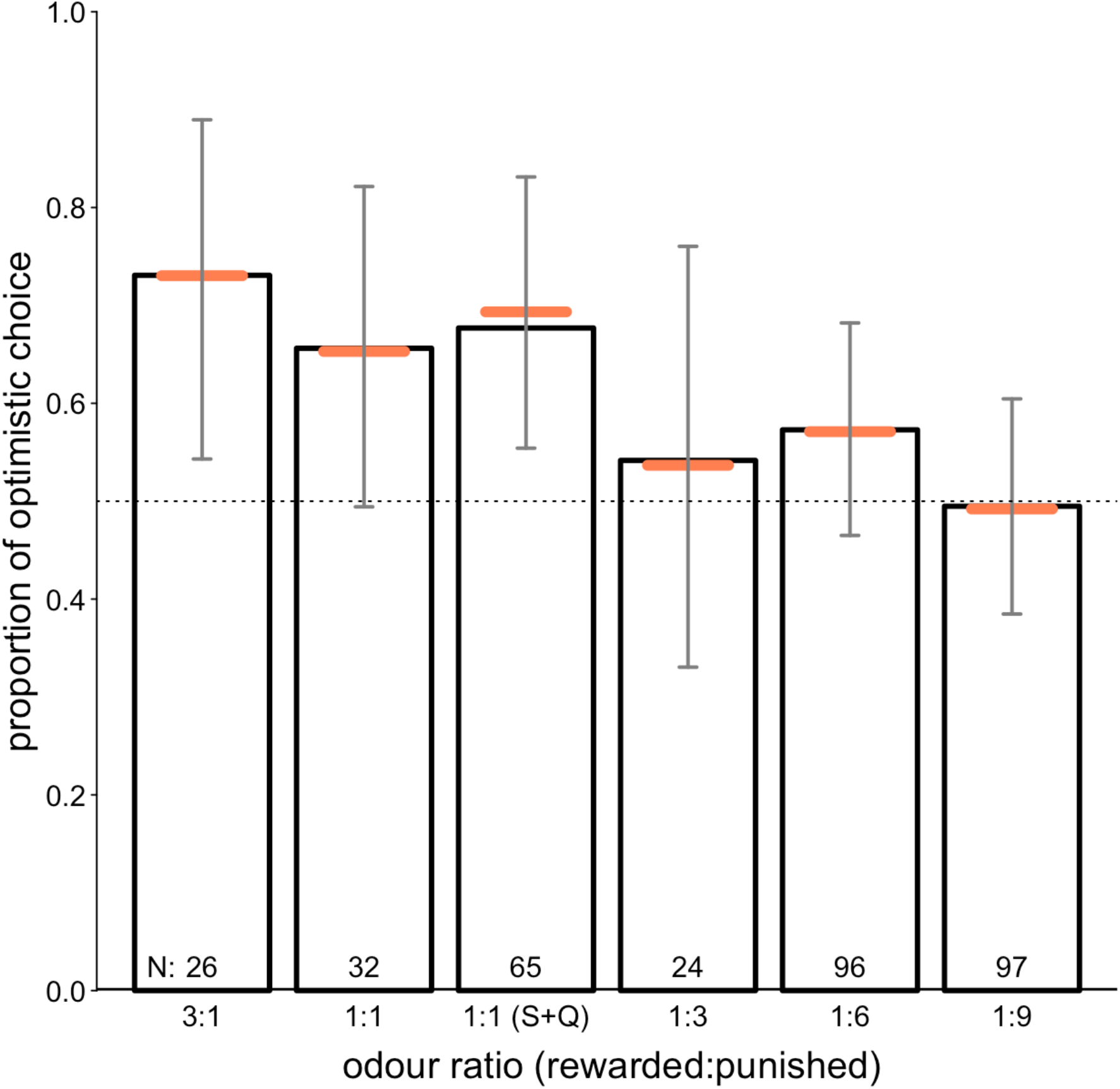
Proportion of optimistic choices (decision for the mixed-odour arm over an unscented arm) for each odour ratio (positive:negative). The horizontal orange lines with error bars depict the fitted model and its 95% confidence interval. Reference categories: 3:1 (ratio), left (rewarded side), lemon (rewarded odour). (S+Q) = shock + quinine.

After completing the learning phase, the subject was allowed up a bridge from the nest leading to a Y-maze. On the Y-maze stem, it was confronted with a solvent control or one of the following positive or negative treatments, using either private or social information:

- 1 µl Dichloromethane (solvent control)
- A 3-volt electric shock (negative private information)
- A tiny 0.2µl drop of 1M sucrose solution, as used in Czaczkes et al. (2019) (positive private information)
- 1 µl undecane, the alarm pheromone of *L. niger* (Bergström & Löfqvist, 1970) (negative social information)
- 1 µl trail pheromone of *L. niger* (following von Thienen et al., 2014) (positive social information)

Note that 3V was used in experiment 2, compared to 9V in the previous experiment, as during data collection in experiment 1b we noted that at least one ant seemed to have been damaged by the stronger voltage.

Subsequently, the subject could decide for the unscented or scented (1:6 ratio) arm of the Y-maze. The ant’s decision was recorded when crossing a 2 cm line on one of the Y-maze arms. Subjects could explore the Y-maze arms for 60 seconds while we recorded the time they spent on the respective scented or unscented arm. Finally, we ran a memory probe to control for learning success by transferring the subject to a second Y-maze, with one arm carrying the positively associated odour and the other arm covered with the negatively associated odour. We recorded the subject’s decision when running at least 2 cm on one of the Y-maze arms. Overall, 302 ants were tested in the second experiment (control: n = 60, shock: n = 60, sucrose: n = 60, alarm: n = 61, trail: n = 61).

The trail pheromone for the social/positive condition was harvested from freeze-killed ants’ hindguts - the glandular source of the *L. niger* trail pheromone (Bestmann et al., 1992; von Thienen et al., 2014). Four glands were dissected and subsequently immersed in 1 ml of dichloromethane. The alarm pheromone solution for the social/negative condition consisted of a 0.01 % (Vol/Vol) concentration of undecane (Bergström & Löfqvist, 1970; Lenz et al., 2013) in dichloromethane. The dilution concentration was based on pilot data (description of methods and results can be downloaded from figshare.com, DOI: 10.6084/m9.figshare.19213017) that showed this concentration to be the minimum to reliably provoke an alarm response and is thus ecologically relevant. Pheromones, both trail and alarm, were applied to the paper layers just before placing them on the Y-maze arms.

### Statistical analysis

The complete statistical analysis code for experiments 1 and 2 can be downloaded from figshare.com (DOI: 10.6084/m9.figshare.19144973). The complete dataset used in the analysis is accessible at the same website (DOI: 10.6084/m9.figshare.19144871). For all analysis, we used R version 4.0.5 (R Core Team, 2021). The alpha level for all analyses was set at *p* < 0.05.

As overall tests of the significance of our main predictors and to avoid cryptic multiple testing (Forstmeier & Schielzeth, 2011) we compared the fit of full Generalised linear mixed-effect models with that of the respective reduced models lacking the main predictor but comprising all other terms present in the full models. The comparisons were based on an Chisq (χ^2^) test.

Finally, the statistical models were validated by examining the distribution of scaled residuals with the simulateResiduals function and testing for over- or under-dispersion using the ‘DHARMa’ package (Hartig, 2019).

The overall sample size in experiment 1 was 340, the overall sample size in experiment 2 was 302. All models contained colony ID as a random effect.

#### Experiment 1: Developing a free-running judgement bias task for ants

In order to assess subjects’ optimistic choice when confronted with odour mixtures of different ratios, we fitted a generalized linear mixed-effects model with binomial error distributions (McCullagh & Nelder, 2019) using the glmer function of the R package lme4 (Bates et al., 2015) with the optimizer ‘bobyqa’. Since our hypothesis was that odour ratio (3:1, 1:1, 1:1 (shock + quinine), 1:3, 1:6, 1:9) influenced our subjects’ optimistic choices (binary response: yes/no), we included odour ratio as our main predictor into our model as well as reward odour (lemon/rose) and the rewarded side (left/right) on which the optimistic choice was located. The model formula was: Optimistic decision = ratio + rewarded side + rewarded odour (+ random effect: colonyID) with ratio, rewarded side and rewarded odour dummy coded and centered. Confidence intervals were obtained by using the function bootMer of the package lme4, using 1,000 parametric bootstraps. Tests of the individual fixed effects were derived by using likelihood ratio tests (Barr et al., 2013, R function drop1 with argument ‘test’ set to “Chisq”). We consequently plotted the fitted model with its estimates and confidence intervals for visual inspection of the model results.

To assess whether subjects performed significantly different in the memory probe trials across odour ratios, we carried out a full-reduced-model comparison with the full model consisting of correct choice (binary response: yes/no) as a response variable as well as odour ratio, rewarded side (left/right) and rewarded odour (lemon/rose) as predictors. The full model formula was: Correct choice = odour ratio + rewarded side + rewarded odour (+ random effect: colonyID) while the reduced model lacked odour ratio.

#### Experiment 2: Judgement biases

Here, we tested whether ants’ tendencies to choose optimistically was influenced by our various treatments fitted a generalized linear mixed-effect model with binomial error distribution using the glmer function of the R package lme4 (Bates et al., 2015). Since our hypothesis was that treatment condition (shock, sucrose, alarm, trail; control as a reference) influenced our subjects’ optimistic choices (binary response: yes/no), we included treatment condition as main predictor into our model as well as reward odour (lemon/rose) and the rewarded side (left/right). The full model formula was: Optimistic choice = treatment condition+ rewarded side + rewarded odour (+ random effect: colonyID) while the reduced model lacked treatment condition as a predictor. We fitted another model with Gaussian error distribution with the same formula but with time spent on the scented Y-maze arm as the response variable, using the function lmer. Like in Experiment 1, we assess the subjects’ learning performance by analyzing the memory probe trials across treatment conditions, using a full-reduced-model comparison. The full model consisting of correct choice (binary response: yes/no) as a response variable as well as treatment condition, rewarded side (left/right) and rewarded odour (lemon/rose) as predictors, while the reduced model lacked treatment condition as a predictor.

## Results

The full statistical output can be downloaded from figshare.com (code and output: DOI: 10.6084/m9.figshare.19144973, dataset: DOI: 10.6084/m9.figshare.19144871).

### Experiment 1: Developing a free-running judgement bias task for ants

Ants showed clear optimistic bias, in that they never showed a preference for the odourless arm, even when the ratio of positive:negative odours was heavily skewed towards the bad (see fig. 1, 95% confidence intervals of model predictions act as statistical tests). Adding an electric shock apparatus on the negatively reinforced runway as an unavoidable reinforcement did not alter the subjects’ optimistic bias (mean proportion of choice for odour mixture ± SD: 1:1: 0.66 ± 0.48, 1:1 (shock + quinine): 0.68 ± 0.47). While the 1:9 ratio mixture of positive and negative odours produced the most indifferent decisions, due to experimental error we decided to use the 1:6 ratio to measure ants’ judgement biases in experiment 2. A model summary comparing all ratios to the 3:1 treatment is provided in supplement Table S1.

The ants showed consistently strong learning in the memory probe, regardless of odour ratio previously tested on (fig. 2). Odour ratio was not a significant predictor in our model (full-reduced model comparison: p = 0.126). Average correct responses were above 72 % for subjects previously confronted with each ratio mixture (mean proportion of correct choices ± SD: 3:1: 0.81 ± 0.4, 1:1: 0.84 ± 0.37, 1:1 (shock + quinine): 0.72 ± 0.45, 1:3: 0.92 ± 0.28, 1:6: 0.84 ± 0.37, 1:9: 0.89 ± 0.32).

**Fig. 2:**
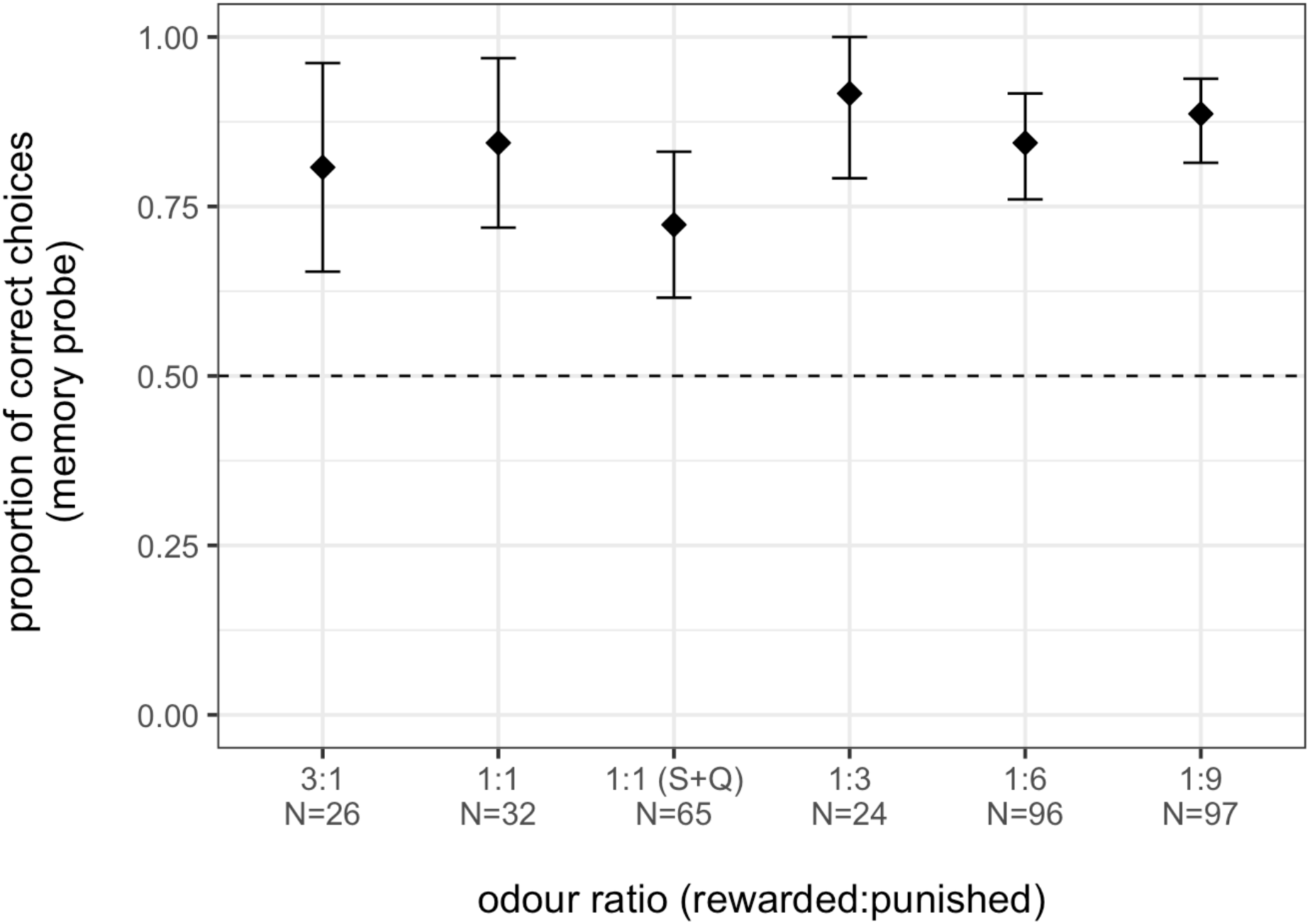
Proportion of correct choices across odour ratios (positive:negative) in the memory probe of experiment 1. Black diamonds show mean proportion per odour ratio, whiskers indicate standard errors. (S+Q) = shock + quinine.

### Experiment 2: Judgement biases

The analysis revealed a clear impact of treatment condition on judgement bias (full-reduced model comparison: p < 0.001), with ants receiving the sucrose treatment choosing significantly more optimistic compared to individuals of the control group. All other treatments, as well as reward odour and scented arm side, did not have a significant impact (table 1, fig. 3), although the shock treatment did elicit a strong trend towards a more negative bias (p = 0.058).

**Table 1:** Results of the optimistic choice model of experiment 2 (estimates, together with standard errors) with control group being the reference category.

| *term* | *estimate* | *standard error* | *z value* | *P* |
| --- | --- | --- | --- | --- |
| *intercept* | -0.491 | 0.336 | -1.464 | 0.143 |
| *shock*<sub>4</sub> | -0.736 | 0.389 | -1.893 | 0.058 |
| *sucrose*<sub>4</sub> | 1.144 | 0.389 | 2.938 | 0.003* |
| *alarm*<sub>4</sub> | -0.117 | 0.368 | -0.318 | 0.750 |
| *trail*<sub>4</sub> | 0.375 | 0.368 | 1.019 | 0.308 |
| *rewarded side*<sub>5</sub> | -0.083 | 0.241 | -0.344 | 0.731 |
| *rewarded odour*<sub>6</sub> | 0.493 | 0.257 | 1.919 | 0.055 |

<sub>4</sub> dummy coded with control being the reference category
<sub>5</sub> dummy coded with left being the reference category
<sub>6</sub> dummy coded with lemon being the reference category

**Fig. 3:**
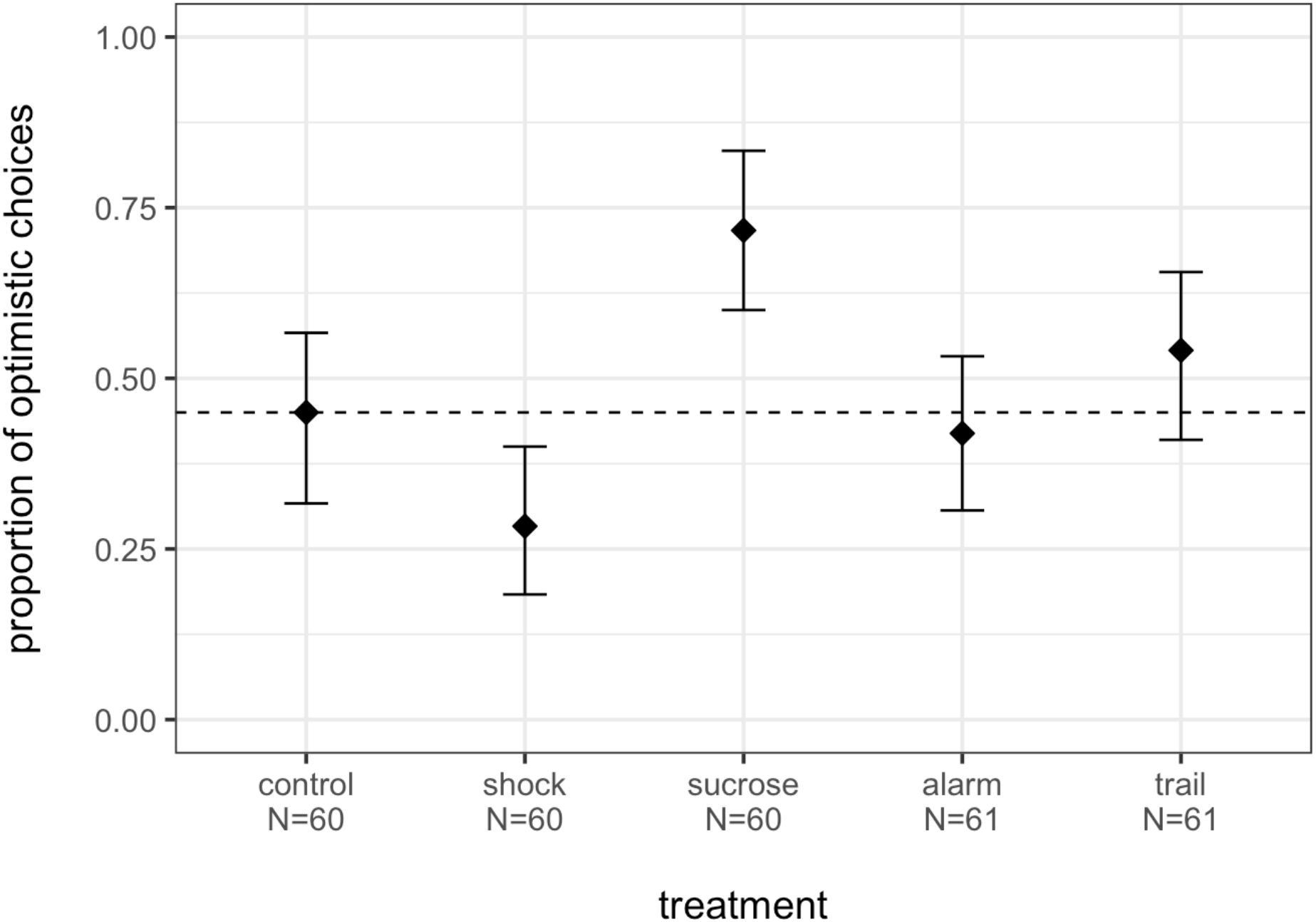
Proportion of optimistic choices across treatment conditions in experiment 2. Black diamonds show mean proportion per treatment condition, whiskers indicate standard errors.

Treatment condition also significantly influenced subsequent performance on the memory probe (full-reduced model comparison: p < 0.001; table S2 in supplementary material, fig. 4). Average correct responses were above 73 % for subjects of all treatment condition but the negative private information treatment. After receiving a shock, ants performed poorly, at around 56 % correct choices, which is significantly below the performance of individuals in the control group (mean proportion of correct choices ± SD: control: 0.733 ± 0.446, sucrose: 0.783 ± 0.415, alarm: 0.871 ± 0.338, trail: 0.869 ± 0.340, shock: mean ± SD: 0.567 ± 0.500, p< 0.05).

**Fig. 4:**
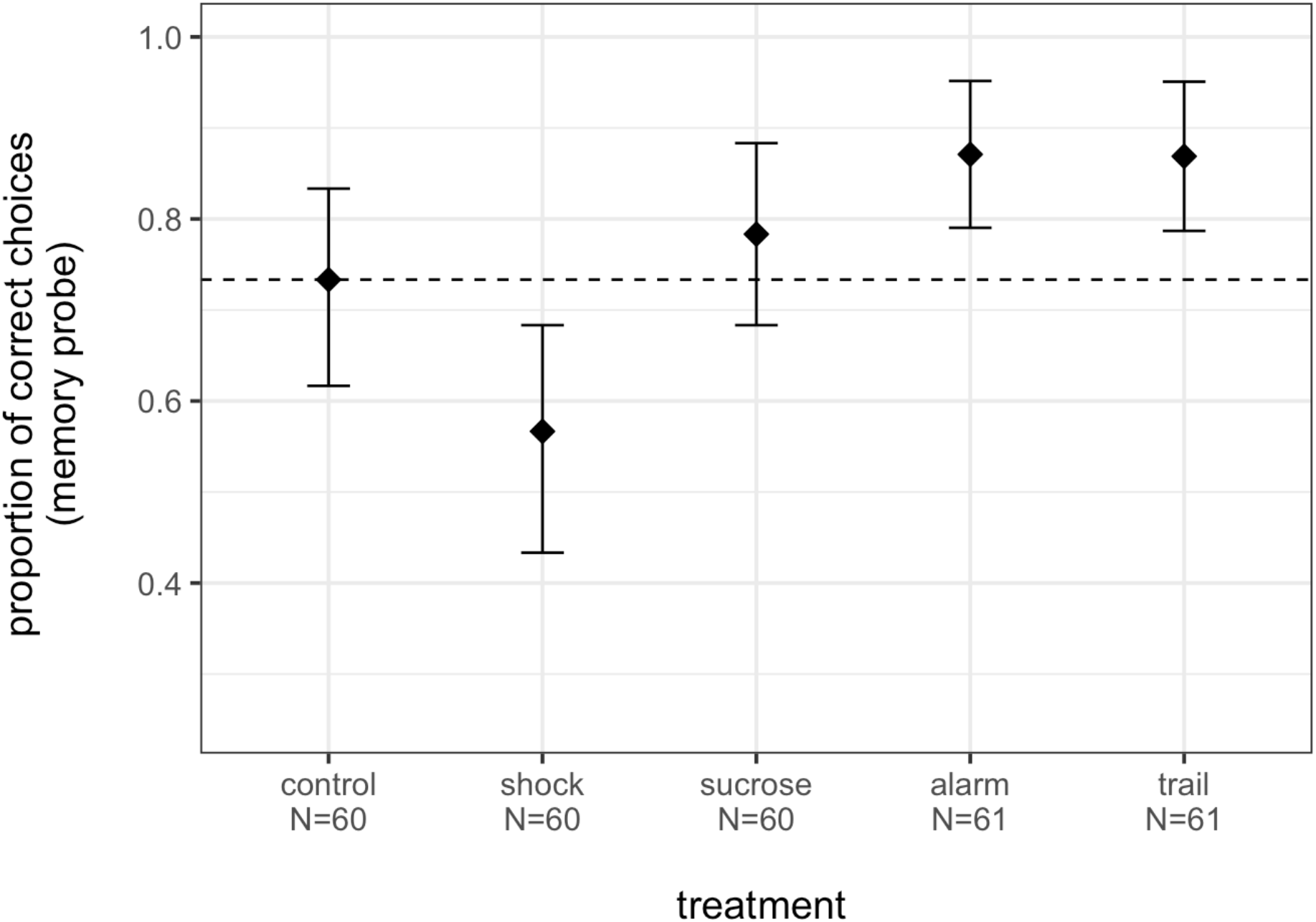
Proportion of correct choices across treatment conditions in the memory probe of experiment 2. Black diamonds show mean proportion per treatment condition, whiskers indicate standard errors.

The full-reduced-model comparison with time spent on scented arm being the response did not reveal a significant effect of treatment condition (full-reduced model comparison: p = 0.672, table S3 in supplementary material, fig. S4 in supplementary material).

## Discussion

Here, we developed an odour-based free-running judgment bias task for insects. Strikingly, subjects chose very optimistically when confronted with an ambiguous 1:1 (positive:negative) ratio. Even given strongly negative-biased ratios of up to 1:9, ants were ambivalent between the mostly-negative odour and no odour at all. Provision of an unexpected reward caused significant positive deviation from baseline judgement bias, and an unexpected negative reinforcement tended to result in a negative judgement bias. Unexpectedly, social information in the form of alarm or trail pheromones did not influence judgement bias.

The ants in our experiment showed a strong positive judgement bias: given an ambiguous cue of 1:1 positive:negative odour, c. 65% ants preferred to follow this over an unscented arm. Even when the negative reinforcement is made stronger and unavoidable by adding a strong (9V) electric shock, a positive judgment bias (68%) remains. This closely matches the findings of Bateson et al., (2011), in which c. 60% of honeybees extended their proboscis in response to 1:1 mix of positive and negative odour. Similarly, while d’Ettorre et al., 2017 did not use a binary choice assay, in their study a larger proportion of ants approached an ambiguous cue in under 100 seconds, as they do towards positively-associated cues, than avoided it, as they do to negative cues. By contrast, results on vertebrate cognitive bias are highly variable, ranging from highly pessimistic (Brilot et al., 2010; Hernandez et al., 2015; J. Novak et al., 2016), through neutral (Hales et al., 2016; Matheson et al., 2008; Murphy et al., 2013; Papciak et al., 2013; Rygula et al., 2012, 2013), to highly optimistic (Brydges et al., 2011, 2012). Comparison of (social) insects and vertebrates is hindered by many vertebrate studies focussing on specific subgroups of individual or treatment conditions. It may well be that social insects are selected to be unusually optimistic: due to reproductive division of labour (Oster & Wilson, 1978) and task partitioning (Ratnieks & Anderson, 1999) individual social insect workers may be under weaker selection to avoid dangers. This is because while solitary animals may risk their entire remaining fitness opportunities, and the efforts already invested into fitness gains, when facing dangers, social insect workers do not, since their colony will continue to function without them. A major goal for cognitive bias research will be to systematically examine bias using ecologically relevant assays over a broad range of study species, in order to understand the selective pressures underlying cognitive bias.

Our findings are in line with previous results in bumblebees (Solvi et al., 2016) that also responded optimistically towards ambiguous cues after receiving a sucrose reward, and with other insects behaving pessimistically after negative experience (Bateson et al., 2011; Deakin et al., 2018; Schlüns et al., 2017). However, caution when interpreting these results as emotions is warranted. The provision of sucrose might not influence emotional states but only increased subjects’ exploratory behaviours, resulting in what seems to be an optimistic response towards the ambiguous stimuli in the judgment bias task (Baracchi et al., 2017). Similarly, it is reasonable to expect recently shocked or shaken animals to have priorities other than feeding – specifically, escape or attack. This would predict a reduction feeding attempts indistinguishable from pessimism in harnessed assays. However, in a free-running assay such as ours this hypothesis would predict random choice, and specifically *not* predict avoidance of the ambiguous cue. Our data suggest that avoidance is in fact taking place (c. 70% avoidance, P = 0.058), in turn suggesting pessimistic judgement bias, as opposed to an escape response.

However, rates of correct choices for the rewarded odour in the memory probe were unusually low for subjects that were previously shocked (Average correct responses: 56%), potentially due to stress-impaired information retrieval (Muth et al., 2015; Schwabe et al., 2012), or a change in motivational state, from foraging to escape. While the balance of parsimony might not be as clearly towards motivational state change as suggested by Baracchi et al., (2017), certainly experiments specifically designed to distinguish emotional bias from motivational state change are sorely needed.

We did not find any indication for emotional contagion, with neither source of social information (trail and alarm pheromone) affecting subjects’ judgment biases compared to control group ants. Whether this is due to a lack of capacity for social transmission of affect in our study species or due to an inefficiency of our methods to elicit emotional or motivational state responses remains unclear. While ants strongly and reliably follow identically prepared pheromone trail solution (von Thienen et al., 2014; Wenig et al., 2021), this has previously been applied as a line, not as a point source of pheromone. While our pilot studies identified a biologically appropriate concentration of undecane, the lack of noticeable alarm response during the alarm treatment in experiment 2 (HK, pers. obs.) implies that a stronger concentration of alarm pheromone may have been able to elicit a change in affect.

The judgement bias task used in this study proved to be a suitable way to assess emotion-like states in ants and holds a great potential for other species which rely primarily on olfaction. Using a choice-based rather than a go/no-go judgment bias task reduced the necessity for inhibitory control (Roelofs et al., 2016). Importantly, it allows the distinction between avoidance and a lack of feeding motivation. Due to its non-invasive nature, the free running assay also kept the impact of the assessment method on the subjects’ natural expressions to a minimum (cf. judgment bias task in a restrained setting, e.g. Bateson et al., (2011)). This method could allow, for example, a broad multi-species comparison of judgement bias, allowing hypotheses about the evolution and ecological relevance of judgement bias to be tested. To briefly give just two examples, colony size may influence judgment bias, with small- colonied species showing less of a positive bias than species with larger colonies (see above). Species from harsh habitats, where food is scarce, may show a stronger positive judgement bias towards food-related ambiguous cues.

Emotions serve to influence information processing and decision-making by directing cognitive resources towards fitness-relevant stimuli, and should therefore be of adaptive value across species (Anderson & Adolphs, 2014; LeDoux, 2012; Paul et al., 2005). Social insects, such as ants and bees, show sophisticated cognitive skills (Chittka, 2017; Czaczkes, 2022a, 2022b; Perry et al., 2017), seem to have the ability to feel pain (Adamo, 2016; Gibbons et al., 2022), have a distinction between “liking” and “wanting” (Huang et al., 2022), and also seem to express emotion-like states (Perry & Baciadonna, 2017 but see Baracchi et al., 2017). Our results therefore feed into the growing discussion on whether insects possess a basic emotion-like system and how they should be included in welfare considerations and regulations (Freelance, 2019; Horvath et al., 2013; Mikhalevich & Powell, 2020). We also hope that our straightforward method for testing judgement bias will accelerate research into judgment bias in invertebrates and open the door to asking broader questions about the evolution and adaptive value of emotion-like states.

## Acknowledgements

We would like to thank Lisa Humbs for her valuable help in data collection as well as all the funding agencies and the University of Regensburg. In addition, we would like to acknowledge Dr. Roger Mundry for his support regarding the statistical analysis.

## Competing interests

We know of no conflict of interest associated with this publication, and there has been no significant financial support for this work that could have influenced its outcome.

## Funding

KW was funded by the Austrian Science Fund (FWF): Doktoratskolleg (DK) program W1262- B29 ‘Cognition and Communication’, and a VDS CoBeNe final fellowship. TJC was funded by a Deutsche Forschungsgemeinschaft (DFG) Emmy Noether grant CZ 237/1-,1 and a <u>DFG</u> Heisenberg grant CZ 237 / 4-1.

## Availability of data, material, and code

The complete statistical analysis code for experiments 1 and 2 can be downloaded from figshare.com (DOI: 10.6084/m9.figshare.19144973). The complete dataset used in the analysis is accessible at the same website (DOI: 10.6084/m9.figshare.19144871).

## Authors’ contributions

KW and TJC designed the experiment, HK and AK performed most parts of the data collection. TJC supervised the project. KW wrote the main manuscript text while all authors discussed the results and contributed to the final manuscript. All authors reviewed the final manuscript.

## Supplementary material

**Table S1:** Results of the optimistic choices model of experiment 1 (estimates, together with standard errors).

| *term* | *estimate* | *standard error* | *z value* | *P* |
| --- | --- | --- | --- | --- |
| *intercept* | 0.872 | 0.477 | 1.829 | 0.067 |
| *1:1<sub>1</sub>* | -0.364 | 0.583 | -0.625 | 0.532 |
| *1:1 (shock + quinine)<sub>1</sub>* | -0.18 | 0.598 | -0.302 | 0.763 |
| *1:3<sub>1</sub>* | -0.85 | 0.606 | -1.401 | 0.161 |
| *1:6<sub>1</sub>* | -0.711 | 0.5 | -1.422 | 0.155 |
| *1:9<sub>1</sub>* | -1.028 | 0.495 | -2.078 | 0.038 |
| *rewarded side<sub>2</sub>* | 0.283 | 0.226 | 1.253 | 0.21 |
| *rewarded odour<sub>3</sub>* | -0.029 | 0.228 | -0.25 | 0.9 |

<sub>1</sub> dummy coded with 3:1 being the reference category
<sub>2</sub> dummy coded with left being the reference category
<sub>3</sub> dummy coded with lemon being the reference category

**Table S2:** Results of the correct choices model of experiment 2 (memory probe; estimates, together with standard errors) with control group being the reference category.

| *term* | *estimate* | *standard error* | *z value* | *P* |
| --- | --- | --- | --- | --- |
| *intercept* | 2.021 | 0.578 | 3.496 | <0.001* |
| *shock*<sub>7</sub> | -0.974 | 0.418 | -2.328 | 0.02* |
| *sucrose*<sub>7</sub> | 0.062 | 0.449 | 0.137 | 0.891 |
| *alarm*<sub>7</sub> | 0.444 | 0.511 | 0.867 | 0.386 |
| *trail*<sub>7</sub> | 0.445 | 0.508 | 0.877 | 0.381 |
| *rewarded side*<sub>8</sub> | -0.395 | 0.3 | -1.316 | 0.188 |
| *reward odour*<sub>9</sub> | -0.019 | 0.338 | -0.056 | 0.955 |

**Table S3:** Results of the time model of experiment 2 (time spent on the scented Y-maze arm; estimates, together with standard errors) with control group being the reference category.

| *term* | *estimate* | *standard error* | *z value* |
| --- | --- | --- | --- |
| *intercept* | 31.649 | 1.502 | 21.065 |
| *shock*<sub>7</sub> | 0.091 | 1.598 | 0.057 |
| *sucrose*<sub>7</sub> | 1.069 | 1.599 | 1.062 |
| *alarm*<sub>7</sub> | 0.222 | 1.611 | 0.138 |
| *trail*<sub>7</sub> | -0.693 | 1.614 | -0.429 |
| *rewarded side*<sub>8</sub> | -2.507 | 1.006 | -2.492 |
| *reward odour*<sub>9</sub> | -0.304 | 1.076 | -0.283 |

<sub>7</sub> dummy coded with control being the reference category
<sub>8</sub> dummy coded with left being the reference category
<sub>9</sub> dummy coded with lemon being the reference category

**Fig. S4:**
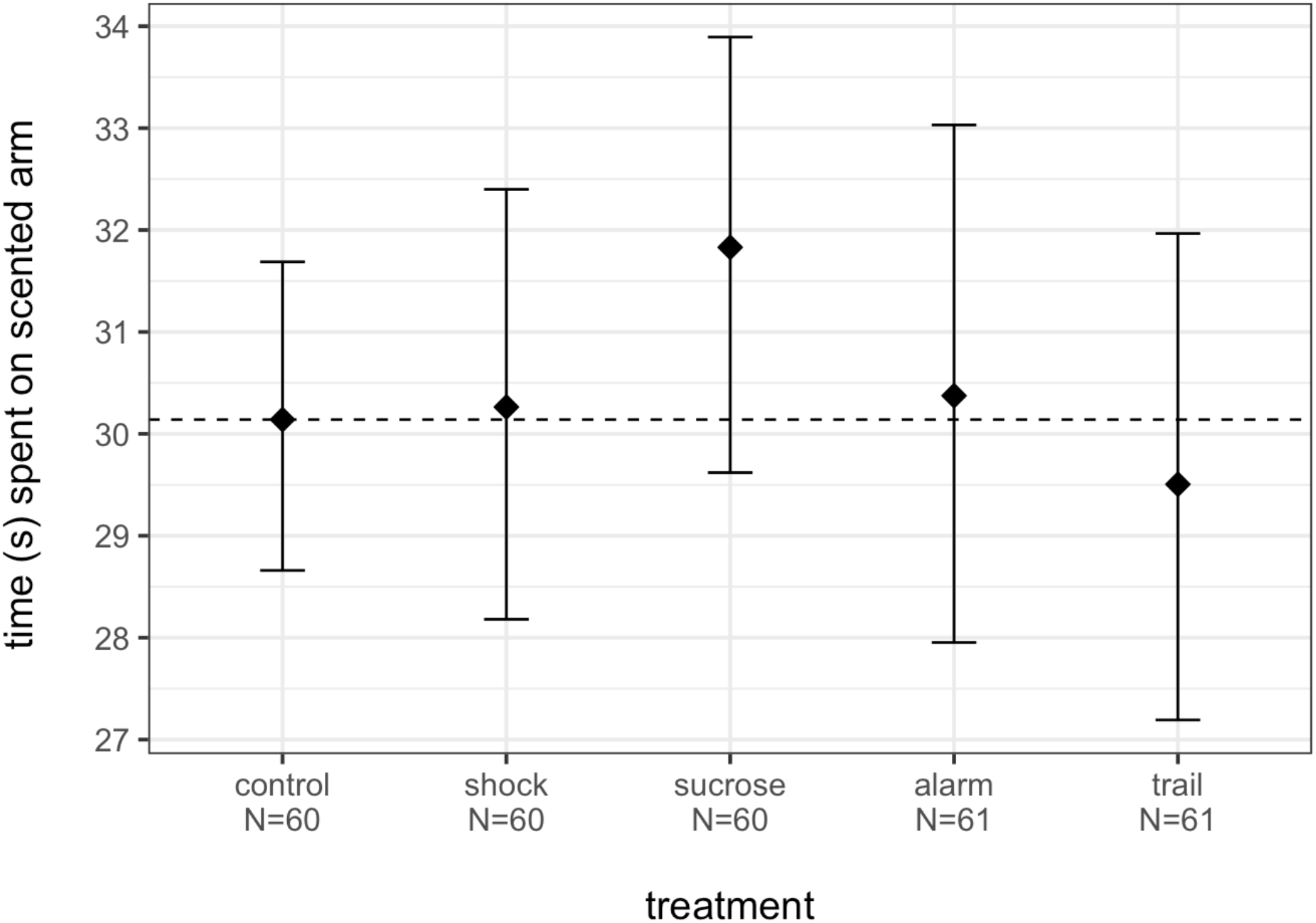
Time spent on the scented Y-maze arm across treatment conditions in experiment 2 (estimates, together with standard errors) with control group being the reference category. Black diamonds show mean proportion per treatment condition, whiskers indicate standard errors.

## Notes

https://figshare.com/articles/presentation/Cognitive_bias_in_Lasius_niger_Undecane_detection/19213017

https://figshare.com/articles/software/Cognitive_biases_in_Lasius_niger_R_script_and_output/19144973

https://figshare.com/articles/dataset/Cognitive_biases_in_Lasius_niger_datasets/19144871

### Competing Interest Statement

The authors have declared no competing interest.

## References

Adamo, S. A. (2016). Do insects feel pain? A question at the intersection of animal behaviour, philosophy and robotics. Animal Behaviour, 118, 75–79. https://doi.org/10.1016/j.anbehav.2016.05.005

Adriaense, Koski S. E., Huber, L., & Lamm, C. (2020). Challenges in the comparative study of empathy and related phenomena in animals. Neuroscience & Biobehavioral Reviews, 112, 62–82.

Adriaense, Martin J. S., Schiestl, M., Lamm, C., & Bugnyar, T. (2019). Negative emotional contagion and cognitive bias in common ravens (Corvus corax). Proceedings of the National Academy of Sciences, 116(23), 11547–11552.

Anderson, D. J., & Adolphs, R. (2014). A Framework for Studying Emotions across Species. Cell, 157(1), 187–200. https://doi.org/10.1016/j.cell.2014.03.003

Baciadonna, L., Duepjan, S., Briefer, E. F., Padilla de la Torre, M., & Nawroth, C. (2018). Looking on the bright side of livestock emotions—The potential of their transmission to promote positive welfare. Frontiers in Veterinary Science, 5, 218.

Baciadonna, L., & McElligott, A. G. (2015). The use of judgement bias to assess welfare in farm livestock.

Baracchi, D., Lihoreau, M., & Giurfa, M. (2017). Do Insects Have Emotions? Some Insights from Bumble Bees. Frontiers in Behavioral Neuroscience, 11, 157. https://doi.org/10.3389/fnbeh.2017.00157

Barr, D. J., Levy, R., Scheepers, C., & Tily, H. J. (2013). Random effects structure for confirmatory hypothesis testing: Keep it maximal. Journal of Memory and Language, 68(3), 255–278.

Bates, D., Mächler, M., Bolker, B., & Walker, S. (2015). Fitting Linear Mixed-Effects Models Using lme4. Journal of Statistical Software, 67(1). https://doi.org/10.18637/jss.v067.i01

Bateson, M., Desire, S., Gartside, S. E., & Wright, G. A. (2011). Agitated Honeybees Exhibit Pessimistic Cognitive Biases. Current Biology, 21(12), 1070–1073. https://doi.org/10.1016/j.cub.2011.05.017

Bateson, M., & Matheson, S. (2007). Performance on a categorisation task suggests that removal of environmental enrichment inducespessimism’in captive European starlings (Sturnus vulgaris). Animal Welfare, 16, 33.

Bateson, M., & Nettle, D. (2015). Development of a cognitive bias methodology for measuring low mood in chimpanzees. PeerJ, 3, e998.

Bergström, G., & Löfqvist, J. (1970). Chemical basis for odour communication in four species of Lasius ants. Journal of Insect Physiology, 16(12), 2353–2375. https://doi.org/10.1016/0022-1910(70)90157-5

Bestmann, H. J., Kern, F., Schäfer, D., & Witschel, M. C. (1992). 3, 4‐dihydroisocoumarins, a new class of ant trail pheromones. Angewandte Chemie International Edition in English, 31(6), 795–796.

Bethell. (2015). A “how-to” guide for designing judgment bias studies to assess captive animal welfare. Journal of Applied Animal Welfare Science, 18(sup1), S18–S42.

Bethell, E., Holmes, A., MacLarnon, A., & Semple, S. (2016). Emotion Evaluation and Response Slowing in a Non-Human Primate: New Directions for Cognitive Bias Measures of Animal Emotion? Behavioral Sciences, 6(1), 2. https://doi.org/10.3390/bs6010002

Bethell Holmes, A., MacLarnon, A., & Semple, S. (2012). Cognitive bias in a non-human primate: Husbandry procedures influence cognitive indicators of psychological well-being in captive rhesus macaques. Animal Welfare, 21(2), 185–195.

Bethell, & Koyama, N. F. (2015). Happy hamsters? Enrichment induces positive judgement bias for mildly (but not truly) ambiguous cues to reward and punishment in Mesocricetus auratus. Royal Society Open Science, 2(7), 140399.

Brilot, B. O., Asher, L., & Bateson, M. (2010). Stereotyping starlings are more ‘pessimistic.’ Animal Cognition, 13(5), 721–731.

Brilot, B. O., Normandale, C. L., Parkin, A., & Bateson, M. (2009). Can we use starlings’ aversion to eyespots as the basis for a novel ‘cognitive bias’ task? Applied Animal Behaviour Science, 118(3–4), 182–190.

Brydges, N. M., Hall, L., Nicolson, R., Holmes, M. C., & Hall, J. (2012). The effects of juvenile stress on anxiety, cognitive bias and decision making in adulthood: A rat model. PLoS One, 7(10), e48143.

Brydges, N. M., Leach, M., Nicol, K., Wright, R., & Bateson, M. (2011). Environmental enrichment induces optimistic cognitive bias in rats. Animal Behaviour, 81(1), 169–175.

Burman, McGowan R., Mendl, M., Norling, Y., Paul, E., Rehn, T., & Keeling, L. (2011). Using judgement bias to measure positive affective state in dogs. Applied Animal Behaviour Science, 132(3–4), 160–168.

Burman, O. H. P., & Mendl, M. T. (2018). A novel task to assess mood congruent memory bias in non-human animals. Journal of Neuroscience Methods, 308, 269–275. https://doi.org/10.1016/j.jneumeth.2018.07.003

Chittka, L. (2017). Bee cognition. Current Biology, 27(19), R1049–R1053.

Clegg, I. L. K., Rödel, H. G., & Delfour, F. (2017). Bottlenose dolphins engaging in more social affiliative behaviour judge ambiguous cues more optimistically. Behavioural Brain Research, 322, 115–122. https://doi.org/10.1016/j.bbr.2017.01.026

Crump, A., Arnott, G., & Bethell, E. (2018). Affect-Driven Attention Biases as Animal Welfare Indicators: Review and Methods. Animals, 8(8), 136. https://doi.org/10.3390/ani8080136

Czaczkes, T. J. (2022a). Advanced cognition in ants. Myrmecological News, 32, 51–64. https://doi.org/10.25849/MYRMECOL.NEWS_032:051

Czaczkes, T. J. (2022b). Advanced cognition in ants. Myrmecological News, 32.

Czaczkes, T. J., Beckwith, J. J., Horsch, A.-L., & Hartig, F. (2019). The multi-dimensional nature of information drives prioritization of private over social information in ants. Proceedings of the Royal Society B-Biological Sciences, 20191136. http://dx.doi.org/10.1098/rspb.2019.1136

d’Ettorre, P., Carere, C., Demora, L., Le Quinquis, P., Signorotti, L., & Bovet, D. (2017). Individual differences in exploratory activity relate to cognitive judgement bias in carpenter ants. Behavioural Processes, 134, 63–69.

Deakin, A., Mendl, M., Browne, W. J., Paul, E. S., & Hodge, J. J. L. (2018). State-dependent judgement bias in Drosophila: Evidence for evolutionarily primitive affective processes. Biology Letters, 14(2), 20170779. https://doi.org/10.1098/rsbl.2017.0779

Detrain, C., Pereira, H., & Fourcassié, V. (2019). Differential responses to chemical cues correlate with task performance in ant foragers. Behavioral Ecology and Sociobiology, 73(8), 107. https://doi.org/10.1007/s00265-019-2717-5

Düpjan, S. (2020). Emotional contagion and its implications for animal welfare. CAB Reviews: Perspectives in Agriculture, Veterinary Science, Nutrition and Natural Resources, 15(046). https://doi.org/10.1079/PAVSNNR202015046

Dyer, F. C. (2002). The biology of the dance language. Annual Review of Entomology, 47(1), 917–949.

Forstmeier, W., & Schielzeth, H. (2011). Cryptic multiple hypotheses testing in linear models: Overestimated effect sizes and the winner’s curse. Behavioral Ecology and Sociobiology, 65(1), 47–55.

Freelance, C. B. (2019). To regulate or not to regulate? The future of animal ethics in experimental research with insects. Science and Engineering Ethics, 25(5), 1339–1355.

Gibbons, M., Versace, E., Crump, A., Baran, B., & Chittka, L. (2022). Motivational trade-offs in bumblebees [Preprint]. Animal Behavior and Cognition. https://doi.org/10.1101/2022.02.04.479111

Hales, C. A., Robinson, E. S., & Houghton, C. J. (2016). Diffusion modelling reveals the decision making processes underlying negative judgement bias in rats. PLoS One, 11(3), e0152592.

Hartig, F. (2019). DHARMa: Residual diagnostics for hierarchical (multi-level/mixed) regression models. R Package Version 0.2, 4.

Hatfield, E., Cacioppo, J. T., & Rapson, R. L. (1993). Emotional contagion. Current Directions in Psychological Science, 2(3), 96–100.

Henry, S., Fureix, C., Rowberry, R., Bateson, M., & Hausberger, M. (2017). Do horses with poor welfare show ‘pessimistic’ cognitive biases? The Science of Nature, 104(1–2), 8. https://doi.org/10.1007/s00114-016-1429-1

Hernandez, C. E., Hinch, G., Lea, J., Ferguson, D., & Lee, C. (2015). Acute stress enhances sensitivity to a highly attractive food reward without affecting judgement bias in laying hens. Applied Animal Behaviour Science, 163, 135–143.

Horvath, K., Angeletti, D., Nascetti, G., & Carere, C. (2013). Invertebrate welfare: An overlooked issue. Annali Dell’Istituto Superiore Di Sanità, 49, 9–17.

Huang, J., Zhang, Z., Feng, W., Zhao, Y., Aldanondo, A., de Brito Sanchez, M. G., Paoli, M., Rolland, A., Li, Z., Nie, H., Lin, Y., Zhang, S., Giurfa, M., & Su, S. (2022). Food wanting is mediated by transient activation of dopaminergic signaling in the honey bee brain. Science, 376(6592), 508–512. https://doi.org/10.1126/science.abn9920

Hunt, J., & Richard, F.-J. (2013). Intracolony vibroacoustic communication in social insects. Insectes Sociaux, 60(4), 403–417.

Josens, R. B., & Roces, F. (2000). Foraging in the ant Camponotus mus: Nectar-intake rate and crop filling depend on colony starvation. Journal of Insect Physiology, 46(7), 1103–1110. https://doi.org/10.1016/S0022-1910(99)00220-6

Knutson, B. (2002). Ultrasonic Vocalizations as Indices of Affective States in Rats. 128(6), 961–977. https://doi.org/10.1037//0033-2909.128.6.961

Lagisz, M., Zidar, J., Nakagawa, S., Neville, V., Sorato, E., Paul, E. S., Bateson, M., Mendl, M., & Løvlie, H. (2020). Optimism, pessimism and judgement bias in animals: A systematic review and meta-analysis. Neuroscience & Biobehavioral Reviews, 118, 3–17. https://doi.org/10.1016/j.neubiorev.2020.07.012

LeDoux, J. (2012). Rethinking the Emotional Brain. Neuron, 73(4), 653–676. https://doi.org/10.1016/j.neuron.2012.02.004

Lenz, E. L., Krasnec, M. O., & Breed, M. D. (2013). Identification of Undecane as an Alarm Pheromone of the Ant Formica argentea. Journal of Insect Behavior, 26(1), 101–108.

Maestripieri, D., Schino, G., Aureli, F., & Troisi, A. (1992). A modest proposal displacement activities as indicator of emotions in primates.pdf. Animal Behaviour, 44(2), 967–979.

Mailleux, A.-C., Detrain, C., & Deneubourg, J.-L. (2006). Starvation drives a threshold triggering communication. Journal of Experimental Biology, 209(21), 4224–4229. https://doi.org/10.1242/jeb.02461

Matheson, S. M., Asher, L., & Bateson, M. (2008). Larger, enriched cages are associated with ‘optimistic’response biases in captive European starlings (Sturnus vulgaris). Applied Animal Behaviour Science, 109(2–4), 374–383.

McCoy, D. E., Schiestl, M., Neilands, P., Hassall, R., Gray, R. D., & Taylor, A. H. (2019). New Caledonian Crows Behave Optimistically after Using Tools. Current Biology, 29(16), 2737-2742.e3. https://doi.org/10.1016/j.cub.2019.06.080

McCullagh, P., & Nelder, J. A. (2019). Generalized linear models. Routledge.

Mendl, M., Brooks, J., Basse, C., Burman, O., Paul, E., Blackwell, E., & Casey, R. (2010). Dogs showing separation-related behaviour exhibit a ‘pessimistic’ cognitive bias. Current Biology, 20(19), R839–R840. https://doi.org/10.1016/j.cub.2010.08.030

Mendl, M., Burman, O. H. P., Parker, R. M. A., & Paul, E. S. (2009). Cognitive bias as an indicator of animal emotion and welfare: Emerging evidence and underlying mechanisms. Applied Animal Behaviour Science, 118(3–4), 161–181. https://doi.org/10.1016/j.applanim.2009.02.023

Mendl, M., Oliver, H. P., & Paul, E. S. (2010). An integrative and functional framework for the study of animal emotion and mood. October, 2895–2904. https://doi.org/10.1098/rspb.2010.0303

Michelsen, A. (2003). Signals and flexibility in the dance communication of honeybees. Journal of Comparative Physiology A, 189(3), 165–174.

Mikhalevich, I., & Powell, R. (2020). Minds without spines: Evolutionarily inclusive animal ethics. Animal Sentience, 29(1).

Moe, R. O., Bakken, M., Kittilsen, S., Kingsley-smith, H., & Spruijt, B. M. (2006). Short communication A note on reward-related behaviour and emotional expressions in farmed silver foxes (Vulpes vulpes)—Basis for a novel tool to study animal welfare. 101, 362–368. https://doi.org/10.1016/j.applanim.2006.02.004

Murphy, E., Nordquist, R. E., Josef, F., & Staay, V. Der. (2014). A review of behavioural methods to study emotion and mood in pigs, Sus scrofa. Applied Animal Behaviour Science, 159, 9–28. https://doi.org/10.1016/j.applanim.2014.08.002

Murphy, E., Nordquist, R. E., & van der Staay, F. J. (2013). Responses of conventional pigs and Göttingen miniature pigs in an active choice judgement bias task. Applied Animal Behaviour Science, 148(1–2), 64–76.

Muth, F., Scampini, A. V., & Leonard, A. S. (2015). The effects of acute stress on learning and memory in bumblebees. Learning and Motivation, 50, 39–47.

Nguyen, H. A. T., Guo, C., & Homberg, J. R. (2020). Cognitive Bias Under Adverse and Rewarding Conditions: A Systematic Review of Rodent Studies. Frontiers in Behavioral Neuroscience, 14, 14. https://doi.org/10.3389/fnbeh.2020.00014

Novak, J., Stojanovski, K., Melotti, L., Reichlin, T. S., Palme, R., & Würbel, H. (2016). Effects of stereotypic behaviour and chronic mild stress on judgement bias in laboratory mice. Applied Animal Behaviour Science, 174, 162–172.

Novak, M. A., Hamel, A. F., Kelly, B. J., Dettmer, A. M., & Meyer, J. S. (2013). Stress, the HPA axis, and nonhuman primate well-being: A review. Applied Animal Behaviour Science, 143(2–4), 135–149. https://doi.org/10.1016/j.applanim.2012.10.012

Orlova, M. (2019). Communication, Pheromones. In C. Starr (Ed.), Encyclopedia of Social Insects (pp. 1–8). Springer International Publishing. https://doi.org/10.1007/978-3-319-90306-4_143-1

Oster, G. F., & Wilson, E. O. (1978). Caste and ecology in the social insects. Princeton University Press.

Papciak, J., Popik, P., Fuchs, E., & Rygula, R. (2013). Chronic psychosocial stress makes rats more ‘pessimistic’in the ambiguous-cue interpretation paradigm. Behavioural Brain Research, 256, 305–310.

Paul, E. S., Harding, E. J., & Mendl, M. (2005). Measuring emotional processes in animals: The utility of a cognitive approach. Neuroscience & Biobehavioral Reviews, 29(3), 469–491. https://doi.org/10.1016/j.neubiorev.2005.01.002

Perry, C. J., & Baciadonna, L. (2017). Studying emotion in invertebrates: What has been done, what can be measured and what they can provide. Journal of Experimental Biology, 220(21), 3856–3868.

Perry, C. J., Barron, A. B., & Chittka, L. (2017). The frontiers of insect cognition. Current Opinion in Behavioral Sciences, 16, 111–118. https://doi.org/10.1016/j.cobeha.2017.05.011

Portha, S., Deneubourg, J.-L., & Detrain, C. (2004). How food type and brood influence foraging decisions of Lasius niger scouts. Animal Behaviour, 68(1), 115–122. https://doi.org/10.1016/j.anbehav.2003.10.016

R Core Team. (2021). R: A language and environment for statistical computing. R Foundation for Statistical Computing (4.0.5). https://www.R-project.org/

Ratnieks, F. L., & Anderson, C. (1999). Task partitioning in insect societies. Insectes Sociaux, 46(2), 95–108.

Reimert, I., Fong, S., Rodenburg, T. B., & Bolhuis, J. E. (2017). Emotional states and emotional contagion in pigs after exposure to a positive and negative treatment. Applied Animal Behaviour Science, 193, 37–42.

Robinson, D. T., Clay-Warner, J., Moore, C. D., Everett, T., Watts, A., Tucker, T. N., & Thai, C. (2012). Toward an unobtrusive measure of emotion during interaction: Thermal imaging techniques. In Biosociology and neurosociology. Emerald Group Publishing Limited.

Roelofs, S., Boleij, H., Nordquist, R. E., & van der Staay, F. J. (2016). Making Decisions under Ambiguity: Judgment Bias Tasks for Assessing Emotional State in Animals. Frontiers in Behavioral Neuroscience, 10. https://doi.org/10.3389/fnbeh.2016.00119

Rygula, R., Papciak, J., & Popik, P. (2013). Trait pessimism predicts vulnerability to stress-induced anhedonia in rats. Neuropsychopharmacology, 38(11), 2188–2196.

Rygula, R., Pluta, H., & Popik, P. (2012). Laughing rats are optimistic. PLoS One, 7(12), e51959.

Saito, Y., Yuki, S., Seki, Y., Kagawa, H., & Okanoya, K. (2016). Cognitive bias in rats evoked by ultrasonic vocalizations suggests emotional contagion. Behavioural Processes, 132, 5–11. https://doi.org/10.1016/j.beproc.2016.08.005

Schlüns, H., Welling, H., Federici, J. R., & Lewejohann, L. (2017). The glass is not yet half empty: Agitation but not Varroa treatment causes cognitive bias in honey bees. Animal Cognition, 20(2), 233–241.

Schwabe, L., Joëls, M., Roozendaal, B., Wolf, O. T., & Oitzl, M. S. (2012). Stress effects on memory: An update and integration. Neuroscience & Biobehavioral Reviews, 36(7), 1740–1749.

Solvi, C., Baciadonna, L., & Chittka, L. (2016). Unexpected rewards induce dopamine-dependent positive emotion–like state changes in bumblebees. Science, 353(6307), 1529–1531.

Stroeymeyt, N., Grasse, A. V., Crespi, A., Mersch, D. P., Cremer, S., & Keller, L. (2018). Social network plasticity decreases disease transmission in a eusocial insect. Science, 362(6417), 941–945. https://doi.org/10.1126/science.aat4793

Tan, S. L. T. (2017). Cognitive bias as an indicator of emotional state and welfare in captive zebrafish.

Von Borell, E., Langbein, J., Després, G., Hansen, S., Leterrier, C., Marchant-Forde, J., Marchant-Forde, R., Minero, M., Mohr, E., & Prunier, A. (2007). Heart rate variability as a measure of autonomic regulation of cardiac activity for assessing stress and welfare in farm animals—A review. Physiology & Behavior, 92(3), 293–316.

von Thienen, W., Metzler, D., Choe, D.-H., & Witte, V. (2014). Pheromone communication in ants: A detailed analysis of concentration-dependent decisions in three species. Behavioral Ecology and Sociobiology, 68(10), 1611–1627. https://doi.org/10.1007/s00265-014-1770-3

Wenig, K., Bach, R., & Czaczkes, T. J. (2021). Hard limits to cognitive flexibility: Ants can learn to ignore but not avoid pheromone trails. Journal of Experimental Biology, 224(11), jeb242454. https://doi.org/10.1242/jeb.242454

Zupan, M., Rehn, T., De Oliveira, D., & Keeling, L. J. (2016). Promoting positive states: The effect of early human handling on play and exploratory behaviour in pigs. Animal, 10(1), 135–141. https://doi.org/10.1017/S1751731115001743

